# Lignocellulose Degradation in Bacteria and Fungi for Biomass Conversion

**DOI:** 10.1101/2024.11.06.622210

**Authors:** Kuan-Ting Hsin, HueyTyng Lee, Ying-Chung Jimmy Lin, Pao-Yang Chen

**Author notes:** These authors contributed equally (Kuan-Ting Hsin, HueyTyng Lee).

## Abstract

Lignocellulose biomass is one of the most abundant resources for sustainable biofuels. However, scaling up the biomass-to-biofuels conversion process for widespread usage is still pending. Bottlenecks during the process of enzymatic hydrolysis are the high cost of enzymes and the labor-intensive need for substrate-dependent enzyme mixtures. Current research efforts are therefore targeted at searching for or engineering lignocellulolytic enzymes of high efficiency. One way is to engineer multi-enzyme complexes that mimic the bacterial cellulosomal system, known to increase degradation efficiency up to 50-fold when compared to freely-secreted enzymes. However, these designer cellulosomes are instable and less efficient than wild type cellulosomes. Fungi cellulosomes discovered in recent years have significant differences from bacterial counterparts and hold great potential for industrial applications, both as designer cellulosomes and as additions to the enzymatic repertoire. Up to date, they are only found in a few anaerobic fungi. In this review, we extensively compared the degradation mechanisms in bacteria and fungi, and highlighted the essential gaps in applying these mechanisms in industrial applications. To better understand cellulosomes in microorganisms, we examined their sequences in 66,252 bacterial species and 823 fungal species and identified several bacterial species that are potentially cellulosome-producing. These findings act as a valuable resource in the biomass community for further proteomic and genetic sequence analysis. We also collated the current strategies of bioengineering lignocellulose degradation to suggest concepts that could be favorable for industrial usage.

## INTRODUCTION

Plant biomass is a resource of renewable energy that is made up of byproducts of agriculture, forestry and the food supply chain. Some examples include wheat straw (1), sugarcane bagasse (2), cereal husk (3) and tree bark (4). With its abundant availability and regenerative nature, plant biomass is therefore a promising candidate to replace fossil fuels as an energy source. The plant biomass resources on earth is estimated to accumulate to about 1.8 trillion tons per year (5) which can be converted to cover more than 80 times of global annual energy (6).

However, this resource is very much under-exploited. Biomass energy only made up of about 5% of total energy production globally (7). The main bottleneck is that multiple components in the entire process of converting plant biomass to biofuel is not cost effective, with no sustainable alternatives available as of now. Take wood biomass as an example, the current optimized steps are pre-treatment to reduce lignocellulose cell wall recalcitrance, followed by enzymatic depolymerization and fermentation of simple sugars to biofuels. There are multiple limitations throughout the process, for example mechanical pre-treatment requires high energy consumption (8), while chemical pre-treatment produces hazardous wastes and require additional neutralization steps (9). Depolymerization and fermentation require enzymes from microorganisms, and therefore are limited by issues such as substrate-specificity, the need for multiple microorganisms to complete the process and complicated product recovery process (10).

One of the ongoing research focus is to improve the conversion efficiency during enzymatic depolymerization. Through increasing the characterization of species and corresponding enzymes that are capable to degrade lignocellulose, the optimization of enzymatic cocktails is constantly being updated (11). This also contributes to designing highly efficient artificial enzyme complexes that are inspired by the cellulose degradation systems found in anaerobic bacteria (12). Characterization of new species is recently enhanced by genomic and bioinformatic advancement, such as the increasing discoveries of heat tolerant and gut cellulolytic microorganisms (13).

Here we review what is known about the natural lignocellulose degradation mechanisms in bacteria and fungi, with the intention to highlight characteristics that can potentially solve bottlenecks in upscaling enzymatic depolymerization of biomass. We summarise current and potential solutions, and introduce prospective species that can be exploited in industrial applications.

## CELLULOSE AND HEMICELLULOSE DEGRADATION

Lignocellulosic biomass consists of three main components, cellulose (60%), hemicellulose (17-32%) and lignin (10-25%) (14–16). Cellulose is composed of a linear polymer chain of 7,000-15,000 glucoses connected by ß-1-4 glycosidic bonds in cross-linked structure which prevents enzyme access (17), forming one of the most stable polymers on earth. The hydrolysis of cellulose can be achieved through oxidative cleavage by at least six types of enzymes. Hemicellulose plays a role of connecting the cell wall skeleton to the cellulose network. It consists of polymers of 5 or 6 carbon sugars linked by glycosidic bonds (16). It has a more complicated structure than cellulose but lower recalcitrance. A diverse set of enzymes are required to fully hydrolyze hemicellulose, including backbone-cleaving enzymes and ancillary enzymes that attack the functional groups. Here we discuss the degradation mechanisms of cellulose and hemicellulose in bacteria and fungi, respectively.

### Cellulose and hemicellulose degradation in bacteria

Cellulose and hemicellulose degradation bacteria can be found in the environment as well as in gastrointestinal system of ruminant animals (18, 19). Both aerobic and anaerobic bacteria are capable to degrade cellulose and hemicellulose. Enzymes that are involved in breaking down complex carbohydrates and polysaccharides into smaller products are known as carbohydrate-active enzymes (CAZymes), which include Glycoside Hydrolase (GH), Carbohydrate Esterase (CE) and Auxiliary Activity (AA) families (20). Microorganism representatives along with their set of cellulose degrading enzymes are listed in Table 1.

**Table 1.**
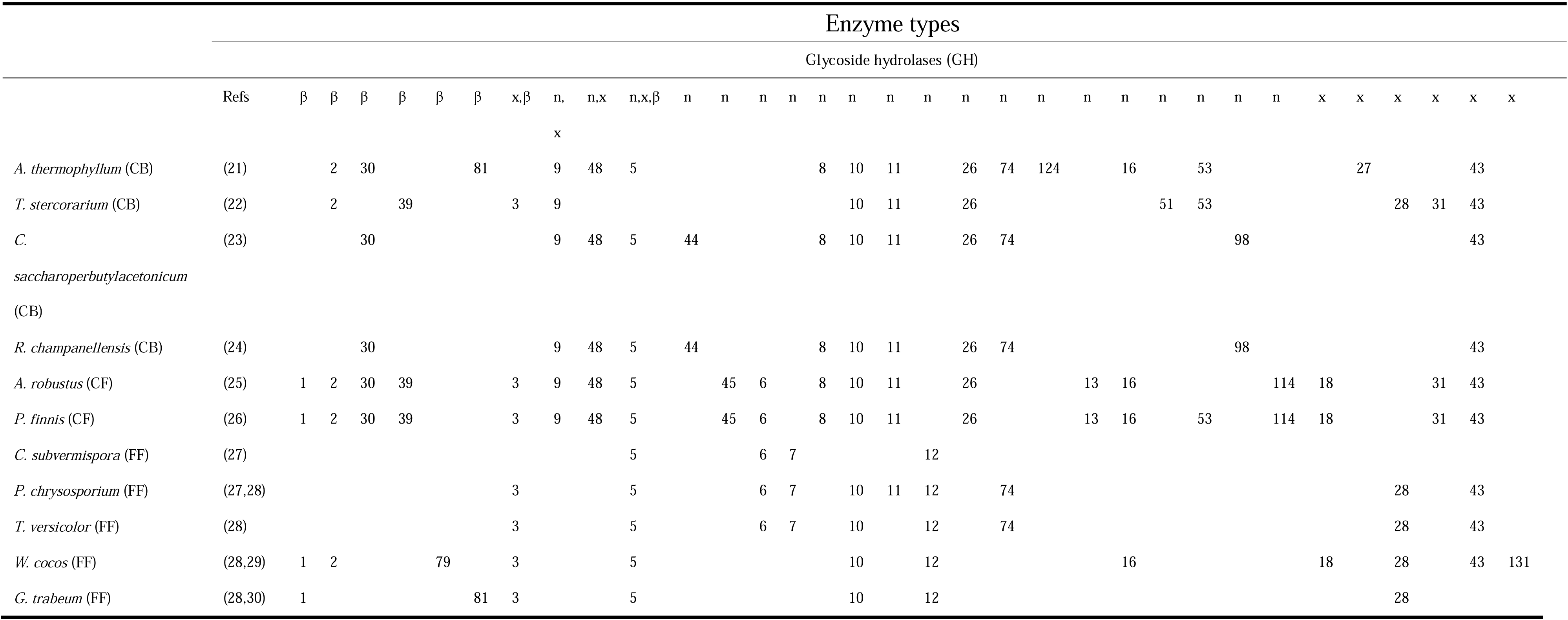
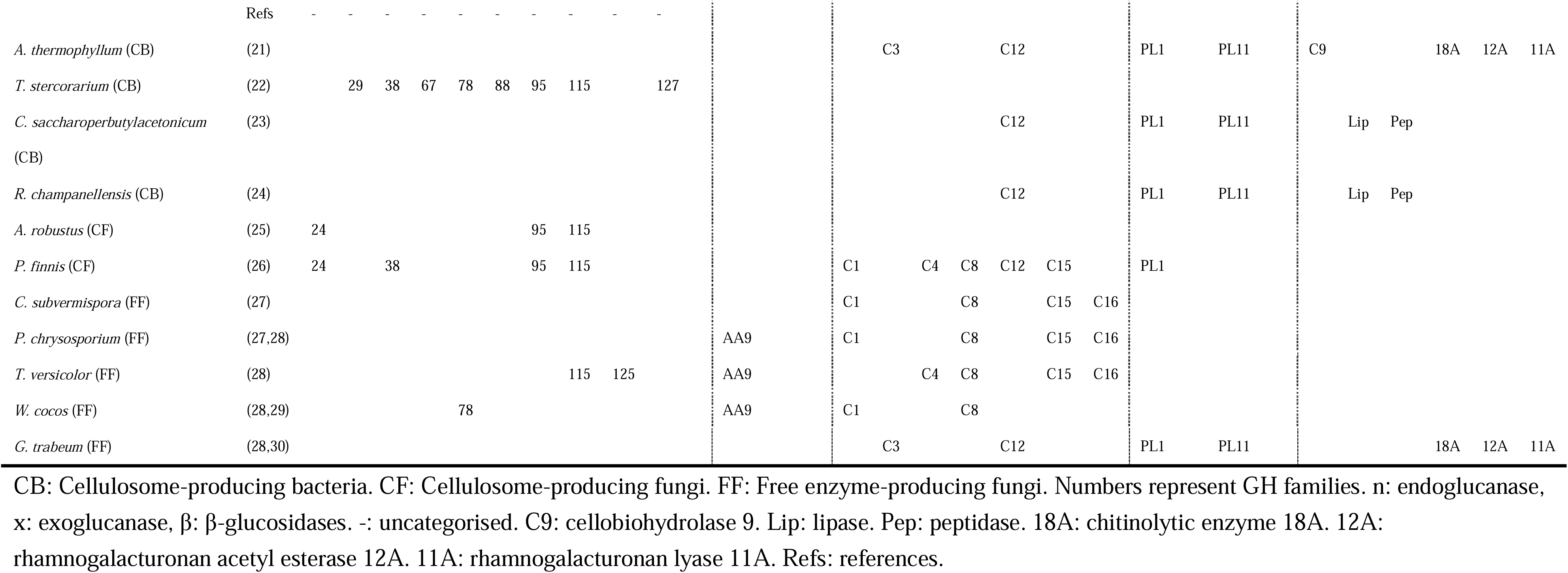
Cellulose-degrading enzymes identified from microorganism representatives.

There are three types of GH families characterized based on the chemical bond cleavage location and the targets to release glucose monomers (16, 19, 21):

1. GH5 to GH12, GH26, GH44, GH45, GH48, GH51, GH74 and GH124 are regarded as endoglucanases due to their ability to cleave 1-4 glycosidic linkage between glucoses randomly (22). Endoglucanases that can act in two types of hydrolysis mechanisms, “retaining” and “inverting” (23).
2. Exoglucanases, like GH3, GH43 and few members from other GH families, are known to act on cellulose terminals to release cellobiose (24).
3. The released cellobiose is then degraded by the β-glucosidases, like GH1-GH3, GH5, GH30, GH39 and GH116 members, which remove the non-reducing end glucosyl residues from saccharides and glycosides to produce glucose (25, 26).

As hemicellulose are made up of xylans, mannans, and xyloglucans, multiple types of endoenzymes are involved in its degradation. To degrade xylan, the xylanase endoenzymes, such as those from GH10, function by hydrolyzing the β-D-xylopyranosyl bonds to release xylobiose, xylooligosaccharides, and xylose (27). As for the mannans, the β-mannanase, β-mannosidase, acetyl mannan esterase, β-glucosidase and α-galactosidase belonging to GH1, GH2, GH3, GH5, GH26, GH27 and GH113 (28) are involved. The GH74, GH29 and GH95 members exhibit the xyloglucan specific endo β 1,4 glucanase activity to degrade the backbone chain of xyloglucan (29).

CEs degrade the ester linkage to release acyl or alkyl groups from carbohydrates, while the AA family consists of copper-based enzymes that cleave the crystalline polysaccharide chain of cellulose via direct oxidative reaction, such as lytic polysaccharide monooxygenases (LPMOs) (30).

### Cellulose and hemicellulose degradation in fungi

Wood decay fungi are fungi that penetrate tree defenses to inhabit their tissues. They utilize the free enzyme mechanism to degrade lignocellulose and are mainly identified within Ascomycetes and Basidiomycetes (31). These fungi are divided into brown-rot, white-rot, and soft-rot fungi based on their ability to decompose different components of wood (32). Brown-rot fungi and soft-rot fungi mainly decay cellulose and hemicellulose, leaving lignin mostly untouched. The accumulation of undecayed lignin in the plant cell wall’s middle lamella makes the wood appear brown and blocky. On the other hand, white-rot fungi selectively decay lignin or simultaneously decay lignin, cellulose and hemicellulose, leaving fibrous and spongy products. By breaking down wood, these fungi play a vital role in the ecosystem, contributing to carbon cycling.

Similar to bacteria, the homologues of cellulose and hemicellulose degrading enzymes like GHs, CEs and AAs are identified from fungi genomic analysis (33, 34). Enzymes like endoglucanases (GH10 and GH11), exoglucanase (GH43), β-glucosidases (GH1 and GH2) and multifunction enzymes (GH5 and GH48), are identified in both brown and white-rot fungi, indicating shared functions in these fungi. Furthermore, a genome-wide search revealed that white-rot fungi generally possess a greater variety of GHs compared to brown-rot fungi, suggesting a divergence in the GH family between these two types of fungi (35). A comparative analysis of GHs in brown-rot and white-rot fungi reveals distinct differences. White-rot fungi encode GHs that can degrade crystalline cellulose, such as GH6 and GH7. These enzymes are missing in brown-rot fungi and is hypothesized to be a result of gene loss events (34).

The fungal CEs exhibit similar functions to those found in bacteria, releasing acyl or alkyl groups attached by ester linkage to carbohydrate backbone of cellulose and hemicellulose (36). With advance in genome analysis, newly characterized fungal CE enzymes from *Pichia pastoris* exhibit both acetyl xylan esterases (AXEs) and feruloyl esterases (FAEs) activities, which facilitate the removal of ester-linked functional group on the linear form of cellulose and hemicellulose to increase the degradation rate (37). The AAs found in fungi exhibit similar function in performing a copper-mediated oxidative break of cellulose at the C1 and/or C4 position of the cellulose chain (38).

Soft-rot fungi composed of a variety of Ascomycota fungi, which are able to live in humid conditions that are unsuitable for brown and white-rot fungi (32). The enzymatic activity of soft-rot fungi is less understood, what is known is that these fungi exhibit an upregulation of genes related to pectinolysis, cellulolysis, and iron acquisition during wood decomposition (39).

Low cellulose recalcitrance mutants, either occurring naturally (40) or genetically engineered (41, 42) had been used to better understand the mechanisms of microfibrils formation. Low recalcitrance is caused by disordered cellulose domains that are shown to be more susceptible to both chemical pre-treatment and enzymatic hydrolysis (43). These disordered domains, termed amorphous cellulose, are the initial targets of “retaining” type endoglucanases (43, 44). Endoglucanases of certain fungal species had been shown to be more effective in targeting amorphous domains, most likely because fungi species have more “retaining” type endoglucanases such as GH7, GH5 and GH12 (45) than the bacterial species.

## LIGNIN DEGRADATION

Lignin is the main source of recalcitrance for biomass (46). It is a network of heterogeneous, alkyl-aromatic polymer that contributes to cell wall integrity and forms an outer barrier for degrading enzymes to access cellulose and hemicellulose (47). The lignin degradation involves the breaking down of alkyl-aromatic polymers into simpler compounds. This process occurs in two steps, depolymerization and aromatic ring cleavage. Depolymerization occurs via the oxidative degradation mechanism, while the aromatic ring cleavage step involves long-range electron transfer process and phenolic moieties oxidation (48). Lignin degradation is better understood in fungi than bacteria. Here we review this process in both systems.

### Lignin degradation in fungi

The white-rot fungi are well-known to degrade the lignin. A diverse set of lignin degradation enzymes is found in white-rot fungi, including dye-depolymerizing peroxidases (DyPs), laccase and laccase-like multi-copper oxidases (LCMOs), lignin peroxidases (LiP), manganese-dependent peroxidases (MnP), laccases (LaC), and versatile peroxidases (VP) (49). DyPs belong to a superfamily of heme peroxidases (50), whereas LCMOs belong to the multicopper oxidase superfamily (51). Both DyPs and LCMOs form two major groups of lignin-degrading enzymes in fungi. The LiPs degrade the aromatic compounds and phenolic compounds in lignin with the presence of H_2_O_2_ via oxidation processes (52). The MnPs generate the diffusible Mn^3+^ from enzymes to lignocellulose structure to form a phenolic compound degradation chelate oxalate (52, 53). The VPs are characterized as hybrid enzymes as they possess both manganese oxidation ability and non-phenolic compounds oxidation ability (49). These enzymes are assigned to the AA family now.

Some brown-rot fungi, like *Gloeophyllum trabeum* also exhibit lignin degradation ability (54). Though both brown-rot and white-rot fungi break down lignin by releasing acyl or alkyl groups (78–81), their mechanisms differ significantly (53, 55, 56). Brown-rot fungi use a non-enzymatic mechanism known as the Fenton reaction, which is an energy-saving process and relies on Fe (II) ions and hydrogen peroxide (49, 56). This reaction reduces Fe (III) to Fe (II) in plant cells, producing -OH radicals that depolymerize lignin polymers (49, 56) and break down cellulose and hemicellulose, making them more accessible to other lignocellulose degradation enzymes.

### Host preference in fungi is affected by ability of lignin degradation

The ability to degrade lignin in different types of fungi contributes to their preference of host (57), as the lignocellulose composition ratio is host-specific. By examining the association between brown and white-rot fungi with woody plants distributed across temperate North America and Europe, white rot was found to favor angiosperms, whereas brown rot preferred gymnosperm (58). A total of 51% of white-rot fungi were specialized to angiosperm hosts, 19% were gymnosperm specialists and 30% were generalists. For brown-rot, 27% were angiosperm specialists, 31% were gymnosperm specialists and 42% were generalists. S-lignin, which possesses an additional methoxyl group to create more variable lignin structure, is present in angiosperm wood but absent in gymnosperm (59). Evolutionarily, brown-rot fungi went through a loss of enzymes that break down S-lignin and therefore are less effective in targeting angiosperms compared to white-rot fungi.

In addition to lignin ratio and structure, difference in wood anatomies also contribute to fungi host preferences. The space for water transportation serves as a “tunnel” for fungi growth. As angiosperms and gymnosperms have different characteristics in these “tunnels”, such as cell diameters, tracheid diameters, size of pits on tangential cell walls, fungi growth and their accessibility for wood degradation differ (60).

### Lignin-degrading bacteria

Lignin-degrading enzymes have been identified genetically (53, 61) and isolated from many bacteria species. However, the degradation strategies in bacteria are less understood, and are believed to be generally less efficient compared to fungi, particularly white-rot fungi (62). Bacteria capable of lignin modification and degradation belong to class Actinomycetes which mainly living in the soil (61), as well as Gammaproteobacteria and Alphaproteobacterial (63).

Compared to fungi, only two classes of lignin-degrading enzymes are identified in bacteria, DyPs and LCMOs (64–66). Bacterial DyPs have relatively lower oxidizing power than fungal DyPs, and therefore target lignin polymers of lower molecular weight (67). Bacterial laccases also have lower efficiency than fungi laccases, but can function in a broader range of environmental conditions (68). With better thermal stability and higher optimal pH (65, 69), bacterial laccases are attractive candidates for industrial usage. Another advantage unique to bacterial lignin-degradation is that bacteria is capable of modifying or converting lignin into valuable bioproducts. Instead of rapidly degrading it in order to access cellulose and hemicellulose, bacteria utilize lignin as a substrate. After lignin monomers are converted into phenolic compounds, aromatic ring cleavage will occur for cellular uptake while producing bioproducts that are of industrial interest such as muconic acid and polyhydroxyalkanoates (70). In addition, bacteria are also easier targets for genetic modification than fungi (71), which allow possibilities of enhancing lignin-degrading enzymes.

## CELLULOSOME

The enzymes described above can either be freely-secreted, extracellular cell-associated, or assembled into a multi-enzyme complex known as cellulosome. As shown in Fig 1A, secreted enzymes diffuse as individual catalytic units and function independently to degrade lignocellulose (55). Species in this category have been extensively exploited for commercial purposes. For example, the fungi *Trichoderma reesei* and *Aspergillus niger* are utilized in the production of cellulolytic cocktails (72).

**Fig 1.**
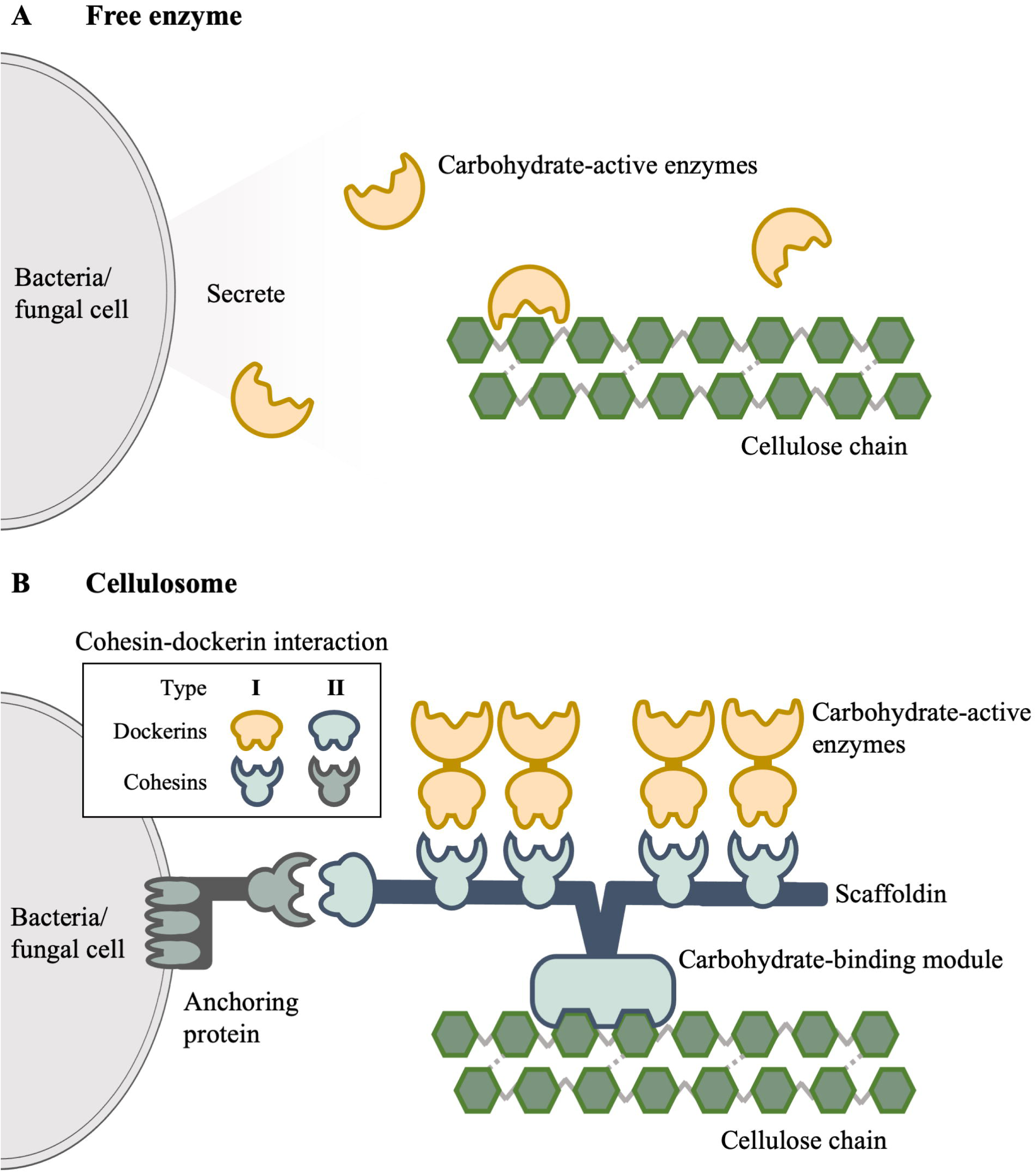
Overview of enzymatic mechanisms of lignocellulose degradation. A Free enzyme mechanism. B Cellulosome.

On the other hand, the cellulosome, first discovered in *A. thermocellum* (73, 74), is a multiprotein complex in which lignocellulolytic enzymes are tethered together for enhanced efficient degradation and systematic hydrolytic activity (75). A simplified cellulosome is depicted in Fig 1B, where it is predominantly situated on the cell wall surface of anaerobic microorganisms with enzymes attached to a large non-catalytic subunit known as scaffoldin. This building-block like structure is assembled through a flexible mechanism involving specialized domains called cohesins and dockerins. The type I cohesin-dockerin interaction allows binding of carbohydrate-active enzymes to scaffoldin (76), while type II interaction allows the scaffoldin to anchor to the cell surface (77). The carbohydrate-binding module (CBM) serves to attach the cellulosome structure to cellulose substrate (16, 77), while the connection of cellulosome to the cell surface is facilitated by surface (S)-layer proteins (78, 79). This multi-enzyme structure is presently exclusive to anaerobic bacteria and fungi that are Clostridia and Neocallimastigomycetes members (75, 77, 80–89) and can be categorized as simple or highly-structured. Simple cellulosome producing bacteria refers to those that produce a single scaffoldin that contains up to nine enzymatic subunits, according to the number of cohesin modules on the scaffoldin, like *Clostridium josui* (90). Highly structured cellulosomes contain several scaffoldins and many enzymes. For example, eight scaffoldins and up to 63 enzymes are identified from *Acetivibrio thermocellum* (79).

There are 22 bacteria and 4 fungi species that were commonly reported in the literature to produce cellulosomes. They are listed in Table 2 with respective references. Class Clostridia consist of anaerobic bacteria that mainly colonize the human gastrointestinal system and soil, whereas Neocallimastigomycetes is a phylum of anaerobic fungi that colonizes the herbivore gastrointestinal system. The strict anaerobic environment and limited energy sources of anaerobic bacteria are two factors that impose selective pressures on them to evolve cellulosomes. This adaptation allows for efficient cellulose degradation to obtain cellular energy (79).

**Table 2.**
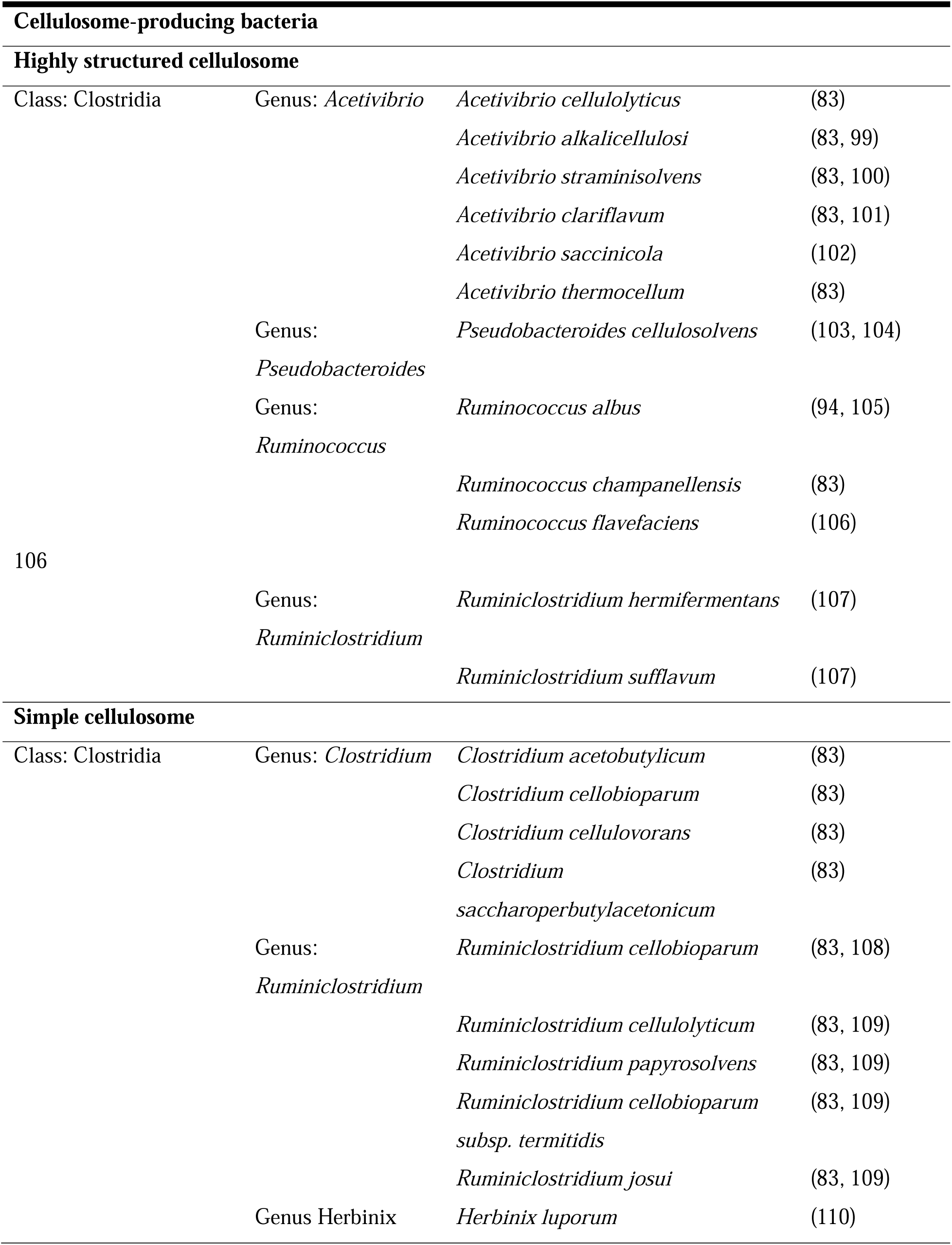
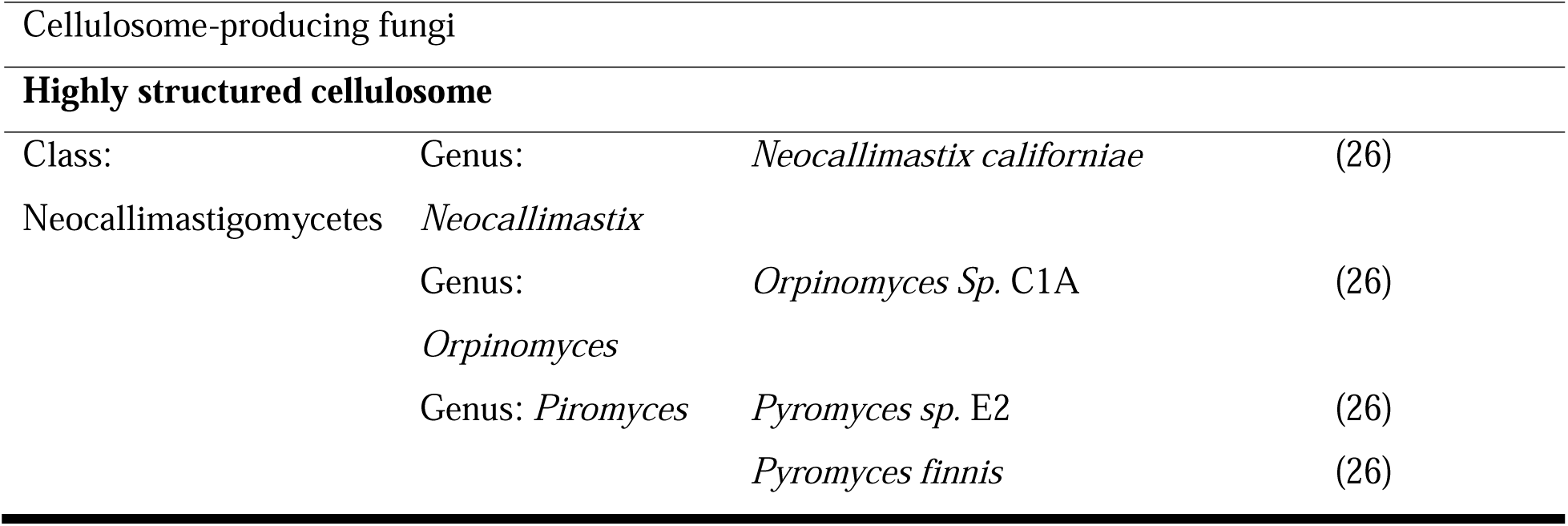
Cellulosome-producing bacteria and fungi.

### Cellulosome in bacteria

The efficiency of bacterial cellulosome has been demonstrated through comparisons of cellulose hydrolysis rate, with *A. thermocellum* achieving a 50-fold increase over free enzymes in *T. ressei* (91) and a 15-fold decrease when *A. thermocellum*’s cellulosome formation is impaired (92). It has been proposed that the spatial enzyme proximity offered by cellulosome structure enhances synergistic interactions between catalytic units (93), while anchoring to the cellulosome increases enzyme stability (79). Another aspect of cellulosomes that remains to be understood is their selective enzyme synthesis based on the substrate. For instance, when grown on differently pretreated yellow poplar and pine-made pulp, *A. thermocellum*’s cellulosomes show varied enzyme compositions (94, 95). Pine pulp, with its higher lignin content, leads to the production of enzymes like xylanase and mannanase for lignin breakdown, while pretreated yellow poplar prompts the synthesis of endoglucanases, exoglucanases, and β-glucosidases (95). Similarly, *Clostridium cellulovorans* produces different sets of enzymes, when grown on cellulose, xylan, galactomannan and pectin (96). These results suggest that enzyme assembly in cellulosomes is influenced by the substrate type.

### Cohesin and dockerin

Components of cellulosomes are held together by high-affinity interactions (97, 98) between *cohesins* and *dockerins*. Three types of cohesin-dockerin interaction were identified so far based on sequence similarity: type I which is responsible for attaching enzymes to scaffoldin, type II which attaches scaffoldin to the cell, and type III which consists of those that are sequentially divergent from another two types (99). Note that this classification is based on bacterial cellulosomes, and the more recently discovered fungal cohesins and dockerins were currently unclassified (100).

Cohesin-dockerin has species- and type-specificity, where cohesins of one species do not interact with dockerins of another species (101), and cohesins of Type I do not interact with dockerins of Type II (79, 102, 103). Within the same species and type, cohesin and dockerin modules have equal affinity and can bind interchangeably (104). The combination of strict species- and type-specificity with module-promiscuity enables accurate assembly of cellulosome with enough heterogeneity of enzyme content, a factor that has been identified to increase degradation efficiency (101).

As cellulosome is an inherently dynamic structure (105), the precise arrangement of the components, including cohesins and dockerins, remains poorly understood in most species. Firstly, the amino acid residues that are responsible for binding specificity and promiscuity as described above is not well understood, as mutational studies only attained conclusive results in limited species (106). Secondly, cohesins and dockerins have been found to play a role in maintaining the plasticity and stability of the whole cellulosome, but the how is not understood. Alternative binding modes have been found in certain types of cohesin (100, 107) which may enable flexibility in the spatial organization of the scaffoldin subunits (108). These binding modes had been hypothesized to be enzymatically regulated by an intramolecular structure located on dockerins. Also, the precise spatial arrangement of enzymatic subunits is found to rely on cohesin-cohesin interactions (105). These findings suggest that cohesins and dockerins are the basis of the regulation of cellulosome stability and plasticity, but quantitative evidence to these hypotheses is still lacking (108).

Thorough understanding of the regulation and organization of cohesin-dockerin interactions can improve the process of lignocellulose degradation, such as through manipulations of the specificity of cohesin-dockerin complexes, as well as enhanced stability and flexibility of artificial designer cellulosomes.

### Cellulosome in fungi

Cellulosomes produced by fungi are less understood, due to the lack of characterized strains and their genetic differences from bacterial cellulosome (109). To date, fungal cellulosomes have been isolated from anaerobic fungi residing in gastrointestinal systems of herbivores (33, 110–112). Despite the early 20^th^-century discovery of fungal cellulosomes in the anaerobic rumen fungus *Neocallimastix frontalis* (113), it was not until recently that their biological characteristics and gene-level components were unveiled (33, 110, 111). This delay was due to the challenges in isolating these organisms, attributed to their anaerobic nature and complex nutritional needs (114). The biological features of fungal cellulosome were initially explored through the herbivore gut-isolated *Piromyces finnis* (110) and the presence of dockerin domains, partial ScaA scaffolding, and GH48 indicate that fungal cellulosomes consist of similar components to bacterial cellulosomes.

In general, there are two key differences in cellulosomes between bacteria and fungi:

1. The fungal dockerin and cohesin domains have no similarity to bacterial counterparts (33). The fungal dockerins form tandem repeats, which are believed to enhance cellulosome binding more than singular, non-repeated domains (115). Contrary to the precise dockerin-cohesin interaction observed in bacteria (93), fungal versions have low species-specificity. Dockerins from one fungal species can attach to cellulosome components of another (116), suggesting the potential for various fungal species within the gut to aid in cellulosome structure formation, thereby boosting its efficiency (33). This cross-binding capability, so far confirmed only in species from the early-diverging fungal phylum Neocallimastigomycota, suggests a possibly widespread conservation of the scaffoldin and dockerin/cohesin system.
2. The second key difference is that fungi with cellulosomes possess the most extensive collection of carbohydrate-active enzymes discovered to date (117). Of all the dockerin-domain-containing proteins in five strains of anaerobic fungi, 13% are GH enzymes that are absent in bacterial cellulosomes (33), therefore contributing enzymatic functions that are unique to fungi. For example, using ß-glucosidase GH3, fungal cellulosome is able to convert cellulose into monosaccharides, instead of the more complex oligosaccharides produced by bacterial cellulosome (118).

Even though the understanding for fungal cellulosome is still based on pieces of evidence, unique features that stood out from bacterial cellulosome may suggest fungal cellulosome to be a potential candidate for industrial applications.

## PRESENCE OF CELLULOSOME PROTEINS IN BACTERIAL AND FUNGAL GENOMES

To discover new species that potentially possess cellulosomes, we scanned all known bacterial and fungal species for the presence of cellulosome proteins, including proteins that form the subunits as well as the enzymes.

### Sequence comparison

First, proteins from bacterial cellulosome were used as baits to search against bacterial and fungal sequences. By using 78 *A. thermocellum* cellulosome proteins as search queries (see Table S1), a sequence similarity search was performed in two databases: 1) the non-redundant database (NCBI nr 20230427) that contains 66,252 bacterial species across 6,823 genera, and 2) the EnsemblFungi database (release 57) that contains 823 fungal species from 419 genera. Query proteins were deemed present in bacterial species if they met alignment criteria of at least 50% identity, an e-value of 10e-2 or lower, and covered at least 70% of the query. For fungi, due to sequence divergence, the criteria were less stringent: a minimum of 50% identity, an e-value of 10e-2 or lower, and at least 50% of query covered. Since only a small amount of *A. thermocellum* cellulosome proteins were present in fungi species, fungal cellulosome proteins were also used as queries: 268 enzymes and 16 potential scaffoldin proteins from *Anaeromyces robustus* were searched in the EnsemblFungi database.

### Novel identification of bacterial species that potentially produce cellulosome

These three sets of searches - *A. thermocellum* proteins in bacterial species, *A. thermocellum* proteins in fungal species and *A. robustus* proteins in fungal species yielded 625 bacterial species, 103 fungal species and 263 fungal species respectively (Table S2 and S3). A total of 22 bacterial species with the highest number of cellulosome proteins found were summarized in Fig 2. Cellulosome protein queries were categorized as originating from both bacteria and fungi (beige; 12 categories), exclusively bacteria (green; 34 categories), or exclusively fungi (blue; 33 categories).

**Fig 2.**
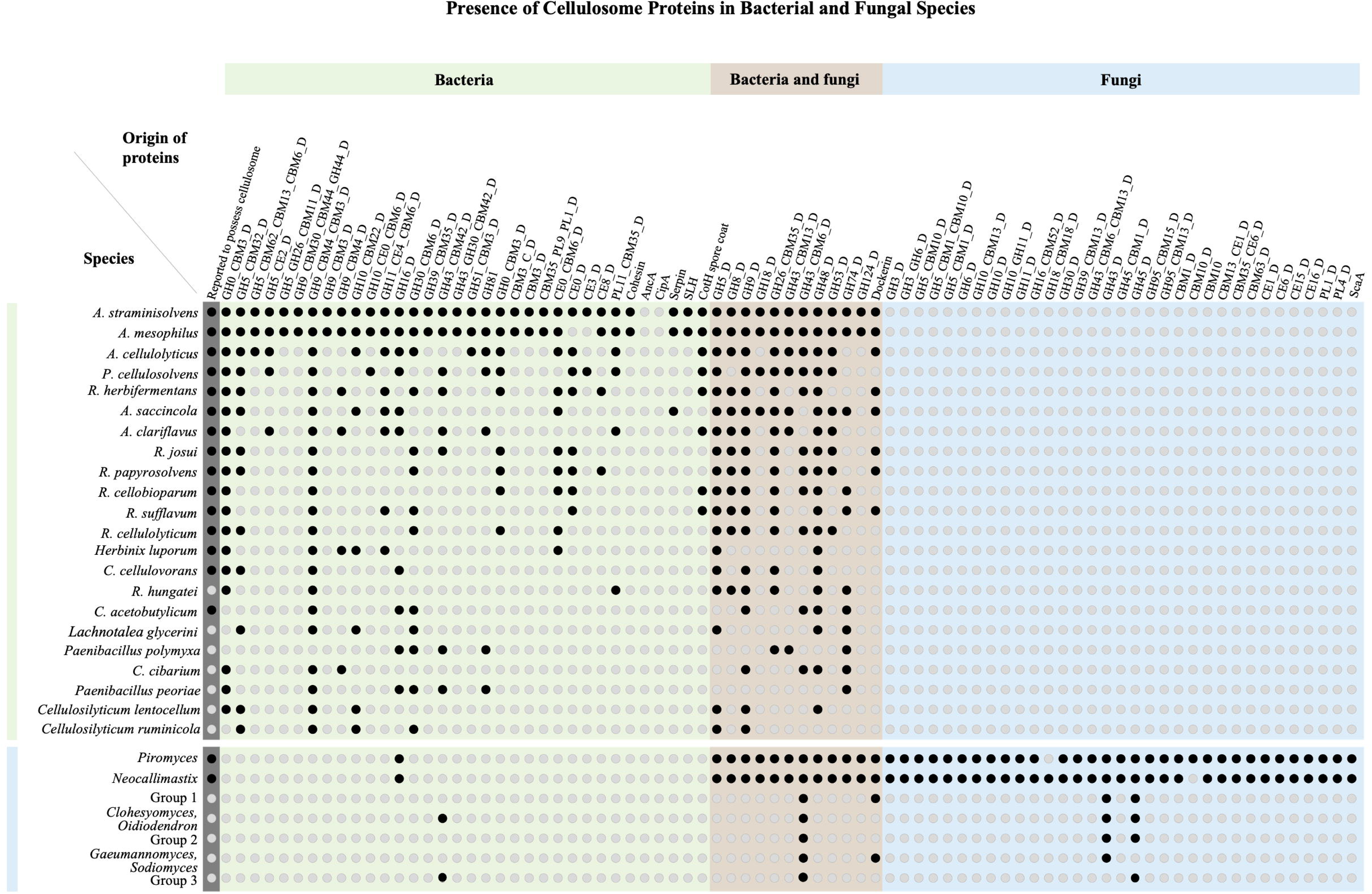
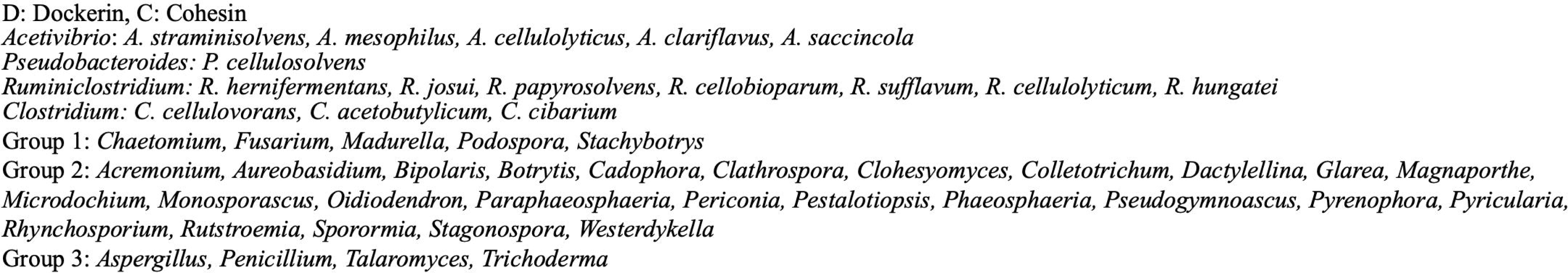
Presence of *Acetivibrio thermocellum* and *Anaeromyces robustus* cellulosome proteins in 17 bacterial species and 41 fungal genera. Protein categories were 1elled as originated from bacteria *(A. thermocellum)* and fungi *(A. robustus)* (beige; 12 categories), from bacteria only (white; 34 categories) and from fungi only (blue; 33 categories).

The results confirm that the sequence alignment approach is effective in identifying the presence of cellulosomes. This is because species with high number of hits have been reported to possess cellulosome. For example, *Acetibivrio* species which are ranked in the top three are known to form complex and elaborated cellulosomal systems (119). Another well-known genus identified is *Ruminiclostridium*, where multiple representatives of this genus are able to produce cellulosomes (120).

There are several species in Fig 2 where the presence of cellulosome is either unconfirmed or has never been explored before. *Ruminiclostridium hungatei* is closely related to other cellulosome-possessing species in the genus such as *R. cellobioparum* and *R. josui* (121), yet whether this species encode cellulosomal genes remained unknown. Large extracellular protein complex had been observed before (122), hinting at the presence of cellulosome. Our results are in alignment with this observation, showing sequence similarities to 8 categories of cellulosomal glycoside hydrolase and polysaccharide lyase. *Cellulosilyticum lentocellum* and *C. ruminicola* are two closely-related species that can degrade cellulose. There is no evidence for cellulosome presence so far, but *C. ruminicola* was found to encode for a subfamily of carbohydrate-binding module (CBM) in its endoglucanase (123). As a component of cellulosome, CBM functions to attach enzymes to substrate. As our results found 7 and 6 cellulosomal protein categories in these species respectively, it is plausible that they could produce cellulosomes. Similarly, *Paenibacillus peoriae* and *P. polymyxa* are two species with high cellulase stability and activity (124, 125), with unknown presence of cellulosome. We found that they encode for 7 cellulosomal protein categories. The possibility for these two species to produce celluosome is high, as one cellulosome-producing species (126) and one CBM-encoding species (127) are identified in the same genus thus far. The presence of cellulosomes in these species can be further confirmed using domain-specific homology analysis and dockerin/cohesin domains binding assays. In addition, enzymes with high sequence similarities to cellulosomal proteins but currently poorly characterized can also be selected for degradation tests with various biomass substrates.

### Non-Neocallimastigomycota fungi probably do not produce cellulosome

As illustrated in Fig 2, the top two genera with the highest number of categories hit are *Piromyces* and *Neocallismastix*, which share the anaerobic phylum with the cellulosome-possessing *A. robustus*. Both *Piromyces* and *Neocallismastix* have 32 out of 33 categories of fungi-only proteins, 12 out of 12 categories of bacteria and fungi proteins, and 1 out of 34 bacteria-only proteins. Since the species from these two genera are known to possess cellulosomes, and the bacterial and fungal cellulosome proteins are known to be sequentially divergent, this result is expected.

For the remaining species, the presence of bacterial and fungal cellulosomal proteins are minimal. At most four categories of proteins are found, three being variations of GH43 and GH45, which are known to be found in fungi that use the free enzyme degradation mechanism, and one being dockerin. The presence of dockerins in non-cellulosomal fungi has been identified previously to function in defensive mechanisms, where these domains form carbohydrate-active toxin complex (128). The presence of dockerins therefore cannot elucidate the presence of cellulosomes, particularly when cellulosomal-specific enzymes GH3 and GH6 are missing in these general.

Among fungi, besides members in the anaerobic phylum Neocallimastigomycota, it appears unlikely that other known species contain cellulosomes. This conclusion aligns with findings from scanning 394 fungal genomes using Hidden Markov Models specific to Neocallimastigomycota scaffoldin, which revealed no sequence similarities (33). It is important to acknowledge that since Neocallimastigomycota is an early-diverged branch, we cannot exclude the possibility that other fungal genera developing highly divergent scaffoldins. Another point to note is that, only few anaerobic fungi species are isolated and sequenced thus far, therefore limiting our search space.

## IMPROVING LIGNOCELLULOSE DEGRADATION FOR INDUSTRIAL USE

Currently, most industrial applications depend on enzymatic cocktails derived from aerobic fungi such as *T. reesei* and *Aspergillus sp.* (129). The significant expense of these mixtures is a major barrier to making biofuels economically viable (130). Moreover, optimizing these mixtures based on the substrate is necessary, as the generic “one-size-fits-all” approach has been proven ineffective (131).

Thus, there are several aspects where bioengineering as well as genomic advancement can help to improve the process:

1. Reducing the concentration of needed enzyme
2. Expanding the enzymatic ability to hydrolyze variable substrates
3. Increasing the enzyme stability in extreme heat or pH conditions that are inevitable during the pretreatment process (130)
4. Reducing the time needed for degradation

Here we discuss three main focus in research development to improve the aforementioned aspects.

### Discovery of new species and enzymes

With more and more species being isolated and sequenced, genomic (such as NCBI) and proteomic (such as CAZy (132)) databases are exponentially increasing in size. Discovering and adding novel lignocellulose-degrading species and enzymes in various ecosystems to the existing repertoire is one of the most valuable effort in improving lignocellulose degradation for industrial use.

One example is the gut anaerobic fungi that remained relatively poorly characterized due to their complex life cycles as well as challenges during recovery and DNA extraction (129, 133). The understanding of cellulosome structures in these fungi has recently improved, but there are more to uncover. Genomic studies of these fungi species improved the understanding of their cellulosome structures, and also showed that their enzyme repertoires offer a competitive edge. They possess four times more carbohydrate-active enzymes than the fungi utilizing free enzyme mechanism (129), with 13% of these enzymes not found in bacterial cellulosomes (33, 75). Moreover, they preferentially degrade more recalcitrant lignocellulose components compared to bacterial cellulosomes (134). This ability from fungi cellulosomal enzymes reduces the necessity for creating unstable, chimeric non-cellulosomal enzymes. Also, a large portion of open reading frames in anaerobic fungal genomes remain unannotated (129) opening the possibility of discovering novel, more efficient enzymes.

The sequence comparison analysis performed in this review is also aimed at contributing to the novel characterization of cellulosome-producing species and carbohydrate-active enzymes. Our results offer list of candidates that could potentially be exploited for lignocellulose degradation in biomass conversion.

### Engineering lignocellulolytic enzymes and cellulosomes

Enzyme engineering is a common practice to modify enzymatic properties through altering the amino acid sequences (135). One of the more popular procedure is through random mutagenesis directed evolutionary, where a target gene in a microorganism is randomly mutated followed by high-throughput screening approaches (136). For example, thermostability is one of the highly desired traits, as besides maintaining reactivity in high temperature, thermostable enzymes can be recycled more efficiently (137). *A. thermocellum* mutants with increased thermostability and reactivity of endoglucanase (138), exoglucanase (139) and ß-glucosidase (140) were successfully generated.

Another strategy for engineering lignocellulolytic enzymes is to mimic the cellulosome structure through constructing chimeric enzymes and form designer cellulosomes. By recombinantly expressing truncated scaffoldins and selected enzyme-dockerin/cohesin chimeras in species that tolerate extreme conditions better (12), designer cellulosomes could be a promising approach for industrial use. Indeed, designer cellulosomes have shown to be 2 to 3 times more effective at breaking down crystalline lignocellulose than free enzymes (98, 141–143).

The main aim of designing cellulosome is to broaden the enzymatic diversity by increasing the enzymes types used. Reconstituted *A. thermocellum* cellulosome with “external” enzymes has been shown to be more effective in degrading crystalline cellulosme (24). Incorporation of non-cellulosomal enzymes such as lytic polysaccharide monooxygenases (LPMOs) and expansins also showed similar effects (144, 145). Cross-species combination of cellulosomal enzymes with dockerin domains has also been shown to have a positive effect on degradation efficiency as glucose production is enhanced when ß-glucosidase from *A. thermocellum* was tagged with the dockerin from *Ruminococcus flavefaciens* (146). The benefit of cross-species combination of enzymes is also proven in fungi, as different types of fungi follow a succession in natural wood decay. The process begins with early colonizers consuming residual sap carbohydrates, followed by intermediate decay by brown-rot and white-rot fungi, and ends with soft-rot fungi thriving in nutrient-rich conditions, ultimately leading to lignin-degraded wood dominated by soft-rot fungi (32, 147). This shows that in order to effectively degrade the lignocellulose, a combination of species is required.

However, the artificial enzyme-dockerin fusion is often instable (148), due to the high species specificity of bacterial cellulosome. Unique traits of fungal cellulosomes might be key to solve this, for example low species-specificity of cohesins and dockerins can enhance compatibility between components; additional C-terminal fusions on dockerins allow capacity for more enzymes to be added to the cellulosome (149).

### Engineering biomass material

Biomass conversion can also be improved through enhancing the material input. Plant materials can be genetically modified to have lower composition of recalcitrant components, such as lower lignin, lower crystalline cellulose and higher amorphous cellulose. There are already successful examples: In woody angiosperm *Populus trichocarpa* CRISPR-Cas9 gene-editing method was applied to generate high syringyl-to-guaiacyl (S/G) ratio mutants with lowered lignin composition (42, 150). In rice, mutants possessed cellulose nanofibrils with a 63% reduction of length, leading to amorphous cellulose chains that serve as initial breakpoints for enzymatic hydrolysis (42). Increased cellulase and β-glucosidase activities were observed, suggesting higher degradation efficiency.

This approach has lower practicality as not all biomass material can be subjected for genetic engineering. Also, for crops materials, yield production is likely to be more prioritized than recalcitrance of cell walls. Nevertheless, research in this direction provides valuable insights into the synthesis and structure of lignocellulose components.

## FUTURE PROSPECTS

The utilization of lignocellulose biomass is a sustainable alternative to fossil carbon resources in producing biofuels to contribute to the circular economy. It is therefore of great interest to develop robust and efficient procedures to degrade lignocellulose. Toward this goal, we thoroughly examined four key aspects. First, we extensively reviewed cellulose, hemicellulose and lignin degradation in bacteria and fungi. Second, we addressed the current understanding of cellulosome, a protein-complex identified in anaerobic bacteria and fungi. Thirdly, we contributed to the attempt of expanding the enzyme repertoire for industrial use by scanning all known bacterial and fungal species for the presence of cellulosome proteins. We identified five bacterial species as potentially cellulosome-producing. In the fourth aspect, we reviewed on directions of research advancements in genomic and bioengineering to improve the process of lignocellulose degradation for industrial use.

By drawing conclusions on lignocellulose degradation systems existing in nature, here a model favorable for industrial usage could be raised. After increasing the surface area of biomass using mechanical pre-treatment, lignin degradation is performed by bacteria AA enzymes. Although the bacterial AA enzymes degrade slower than fungi, they have the advantage of heat tolerance and can produce economical byproducts. The bacterial system is also an easier target for bioengineering, so lignin degradation efficiency can be further enhanced genetically. After lignin is removed, fungal endoglucanase should be used to degrade amorphous cellulose. Once the amorphous cellulose is broken down, designer cellulosomes can be used to hydrolyze the crystallized parts of cellulose and hemicellulose. The designer cellulosome is recommended to be assembled using fungal scaffoldin backbone and cohesin/dockerin modules to capitalize on their cross-binding ability. Enzymes attached on the designer cellulosome should be from mixed species of bacteria and fungi. To ensure that the enzymes work synergistically, the arrangement of the enzymes on the cellulosome should take the natural succession of microorganism communities into consideration. For example, enzymes from sugar fungi should come before intermediate decay fungi and followed by soft rot fungi.

In the future, we anticipate answers to biological questions such as what is the mechanism behind the secretion of a selected choice of substrate-specific synergistic enzymes, how and when is the cellulosome assembled, what selection factors contribute to the repeated evolution of cellulosomal structure in both bacteria and fungi, etc. These insights, together with a wider collection of synthetic biology tools, will help actualize the usage of biomass as a sustainable energy.

## FUNDING

This work was supported by Academia Sinica (NTU-AS Innovative Joint Program: AS-NTU-112-12), and the National Science and Technology Council Taiwan Ministry of Science and Technology (112-2311-B-001-007 and 111-2927-I-001-505 to P.-Y.C.)

## AUTHOR CONTRIBUTIONS

P.Y.C. designed the research. K.T.H and H.T. L and Y.C.L. performed extensive literature review and write the manuscript. H.T. L analyzed the data. P.Y.C. and Y.C.L. edited the manuscript.

## SUPPLEMENTARY MATERIALS

Table S1 A total of 78 *A. thermocellum* and 284 *A. robustus* proteins categorized based on their domains and functions in the cellulosome structure. Table was deposited in Zenodo repository under 10.5281/zenodo.13324669.

Table S2 BLAST results of aligning 78 *A. thermocellum* proteins to the non-redundant database. Filtering criteria: at least 50% percentage identity, e-value of 10e-2 and below, at least 70% of query covered and within the bacteria taxonomy. Table was deposited in Zenodo repository under 10.5281/zenodo.13324669.

Table S3 BLAST results of aligning 78 *A. thermocellum* proteins and 284 *A. robustus* proteins to the EnsemblFungi database. Filtering criteria: at least 50% percentage identity, e-value of 10e-2 and below, and at least 50% of query covered. Table was deposited in Zenodo repository under 10.5281/zenodo.13324669.

## DATA AVAILABILITY

NCBI and JGI protein IDs of *Acetivibrio thermocellum* and *Anaeromyces robustus* used as alignment queries were listed in Table S1.

The two public databases used were non-redundant database (NCBI nr 20230427) and EnsemblFungi database (release 57).

## Notes

### Competing Interest Statement

The authors have declared no competing interest.

